# Direct detection of the myosin super-relaxed state and interacting heads motif in solution

**DOI:** 10.1101/2021.06.29.450383

**Authors:** Sami Chu, Joseph M. Muretta, David D. Thomas

**Affiliations:** Department of Biochemistry, Molecular Biology, and Biophysics, University of Minnesota, Minneapolis, MN 55455

**Keywords:** myosin, fluorescence resonance energy transfer (FRET), structure-function, enzyme structure, cardiac muscle

## Abstract

We have used time-resolved fluorescence resonance energy transfer (TR-FRET) to detect the interacting-heads motif (IHM) of β-cardiac myosin in solution. Evidence for the IHM has been observed by several structural techniques, and it has been proposed to be the structural basis for the super-relaxed state (SRX), a low-ATPase state of myosin that has been observed biochemically in skinned muscle fibers using fluorescent ATP. It has been proposed that the disruption of this state, by mutation or chemical modification, is a major cause of heart disease, so drugs are being developed to stabilize it. The goal of the present study is to determine directly and quantitatively the correlation between the measured fractions of myosin in the IHM state and the SRX state under the same conditions in solution. We used TR-FRET to measure the distance between the two heads of bovine cardiac myosin, and found that there are two distinct populations, one of which is observable by FRET at a center distance of 2.0 nm, and the other is not detected, implying a distance greater than 4 nm. Under the same conditions, we also measured the fraction of heads in the SRX state using fluorescent nucleotide and stopped-flow kinetics. We found that, in the absence of crosslinking, the population of SRX exceeded that of IHM. In particular, the stabilizing effect of mavacamten was much greater on SRX (55% increase) than on IHM (4% increase). We conclude that the SRX and IHM states are related, but they are not identical.

## Introduction

Myosin II is a motor protein that powers the contraction of muscle through large structural changes when its ATPase activity is activated by actin (1). It consists of a dimer of heavy chain (HC) molecules each binding two light chains, the essential light chain (ELC) and regulatory light chain (RLC). In striated muscles (skeletal and cardiac), myosin polymerizes to form bipolar thick filaments through coiled-coil interactions of the C-terminal tail region of the HC. The N-terminal region branches from the tail to form two heads, each containing a globular catalytic domain and a light-chain domain that binds ELC and RLC (Fig. 1A). The present study uses a cleaved two-headed version of myosin, heavy meromyosin (HMM), which includes the S2 portion of the tail that stabilizes the dimer.

**Figure 1.**
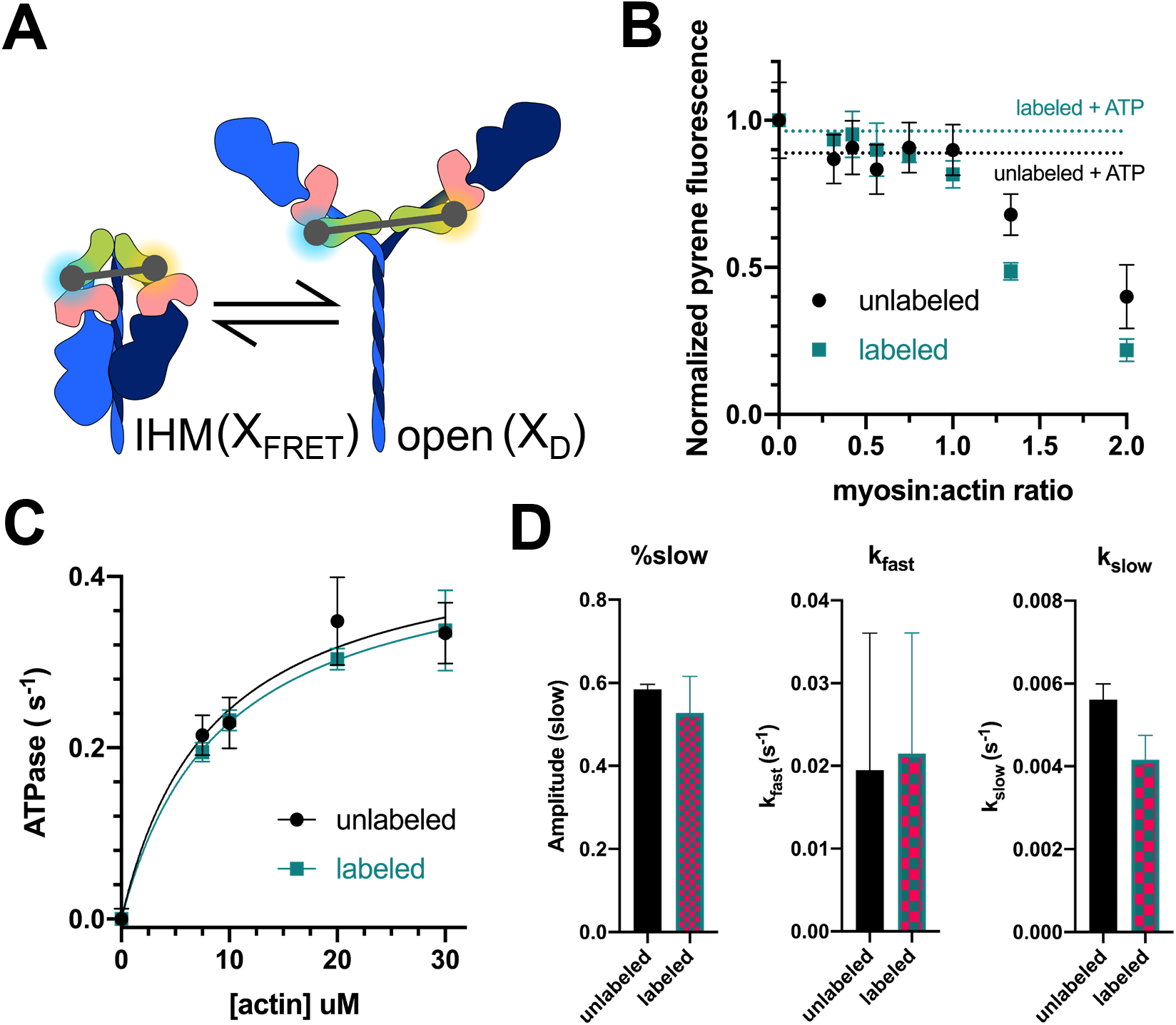
Labels on HMM have no effect on myosin function. (A) Schematic representation of IHM and 6S myosin molecule with FRET labels attached on the RLC. Heavy chain is in blue, with the blocked head of the IHM in dark blue and free head in light blue; ELC is in green, and RLC is in pink. FRET donor (EDANS) is in blue, and FRET acceptor (dabcyl) is orange. (B) Actin-binding function of labeled HMM measured by fluorescence quenching of pyrene-labeled actin. Dotted line indicates fluorescence after the addition of 2mM ATP. (n=4 per condition, p = 0.90 in 2-tailed unpaired t-test). (C) Steady-state ATPase activity of labeled HMM measured by NADH-coupled ATPase activity (n=3 per condition, p = 0.75 in 2-tailed unpaired t-test). (D) Basal ATP turnover measured with and without labels (n=3 per condition).

The regulation of muscle contraction involves primarily the calcium-dependent regulation of the actin-containing thin filament. However, an additional regulatory mechanism has recently come to light, in which myosin regulates itself through autoinhibitory interactions between its two heads. It is proposed that the structural basis for this autoinhibited state is the interacting-heads motif (IHM) (2–5). Evidence for the IHM has been obtained in several isoforms of myosin from several species through electron microscopy (3,5–7), and evidence consistent with it has been observed by x-ray diffraction (8,9) and fluorescence polarization (10,11). In the IHM, the catalytic domains of the myosin dimer interact asymmetrically - the converter domain of the free head blocks the actin-binding region of the blocked head (3). Another region of interest in the IHM is the mesa region of the catalytic domain, which is proposed to interact with the portion of the tail region closest to the catalytic domain (also called subfragment-2 or S2) (12,13) (Fig. 1A). Studies have shown that disease-related mutations in this region decrease the affinity of the catalytic domain for S2, thus destabilizing the IHM (13).

It has been hypothesized that the IHM is the structural basis for the super-relaxed state (SRX), a functional state characterized by extremely slow ATP turnover, 100 times slower than during contraction, and 3-10 times slower than is typical for most of the myosin molecules in relaxed muscle (14). SRX is detected biochemically in single-turnover experiments, measuring the decrease in fluorescence that occurs when fluorescent mant-ATP is hydrolyzed, released from myosin, and displaced by non-fluorescent ATP, resulting in a multiple exponential decay, in which the slowest rate (longest lifetime τ) is assigned to the SRX state (14). This kinetic signature has been observed in isolated myosin molecules and fragments (15). It is generally assumed that the SRX biochemical kinetic state corresponds to the IHM structural state, but this has not been tested directly on the same sample under similar conditions.

To bridge this gap, we have used fluorescence resonance energy transfer (FRET) between the two heads of myosin to detect the IHM structural state directly in solution, and directly compare the fraction of myosin molecules in SRX measured by ATP turnover, under similar conditions. In contrast to intensity-based FRET measurement, time-resolved FRET can resolve multiple distinct populations of protein conformations, each giving rise to a distinct lifetime τ of decay after an exciting pulse. To verify that FRET is detecting the IHM, we used a chemical crosslinker that are known to stabilize the IHM.

To further evaluate the relationship between IHM and SRX, we compared how both the fluorescent nucleotide turnover and the structural FRET measurement were affected by mavacamten, a small molecule that is currently in Phase III clinical trials for treating hypertrophic cardiomyopathy and has been shown to increase the SRX population (15,16). Given the current state of knowledge, we expected that mavacamten would increase FRET by increasing the fraction of myosin in IHM.

## Results

### FRET labels have no effect on myosin function in the presence of actin

Purified bovine cardiac HMM myosin was labeled with EDANS and dabcyl through the exchange of RLC labeled on the C-lobe by sulfhydryl reaction between an exogenous cysteine residue on the protein and maleimide on the label, as described in our previous work (Fig 1A) (15). To minimize the donor-only population, the ratio of donor to acceptor was 1:3, leading to detectable FRET (Fig 2A). Labeling did not significantly affect the actin association and disassociation (Fig 1B), actin-activated ATPase activity (Fig 1C) nor single nucleotide turnover kinetics (Fig 1D).

**Figure 2.**
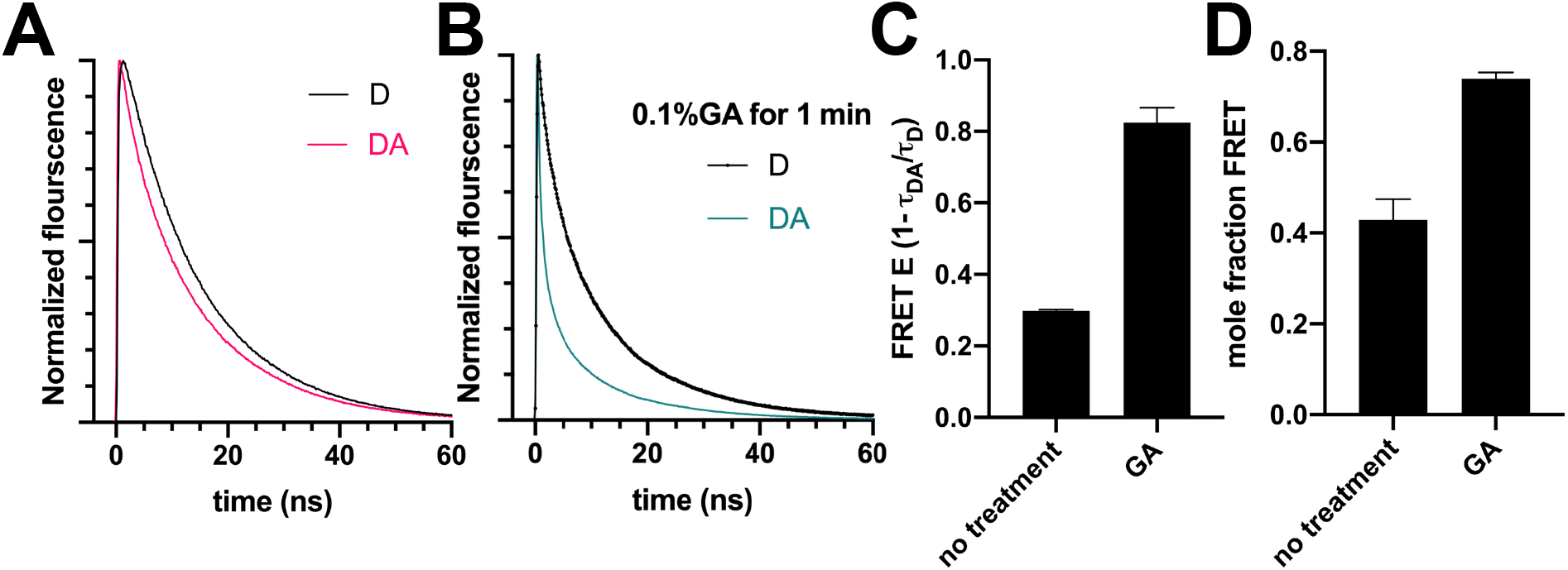
TR-FRET detects the IHM, and treatment with glutaraldehyde increases FRET. (A) Representation waveforms for donor-only (D) and donor-acceptor (DA) samples (B) Representation waveforms for donor-only (D) and donor-acceptor (DA) samples post-glutaraldehyde treatment (C) Changes in %FRET due to glutaraldehyde (n=3 per condition) (D) Changes in mole fraction of population participating in FRET

### Time-resolved FRET detects the closed state of HMM myosin

Time-resolved fluorescence of labeled HMM was best fit by a two-component model: 63% of the donors were unaffected by FRET, having a lifetime identical to the donor-only sample, while the remaining 37% (±5%) showed substantial FRET, fitting to a Gaussian distance distribution having a center of 2.0 nm (±0.02) with a full-width of 1.4 nm (±0.02). We conclude that myosin molecules with the open conformation have a donor-acceptor distance that is too large to detect, and that approximately 40% of the myosin molecules are in the IHM state (Fig 2D).

To verify that we are detecting the IHM, we used a variety of myosin effectors known to either increase the population of IHM or SRX myosin.

### The closed state population increases with the trapping of IHM myosin by crosslinker

To test whether the FRET pair on opposite heads of HMM can measure IHM, labeled HMM was treated with glutaraldehyde, a crosslinker used to trap the IHM in electron microscopy studies (3). We found that glutaraldehyde significantly increased FRET (Fig 2C).

As a control, a donor-only sample was also treated with glutaraldehyde. Glutaraldehyde caused changes in EDANS fluorescence. We used the same two-state model as above to fit a fraction donor-only and a fraction participating in FRET, and found that the fraction participating in FRET increased from 43% (±5%) to 74% (±1%) with glutaraldehyde treatment. 75% is the value predicted if all myosin molecules are participating in FRET, based on the 1:3 donor to acceptor ratio. supporting our conclusion that FRET detects the IHM structure.

### Mavacamten, a stabilizer of the SRX, increases the closed state population less than predicted

Mavacamten is a Phase-III drug used to treat hypertrophic cardiomyopathy. It has been shown to significantly decrease myosin activity, and electron microscopy has shown that cardiac myosin treated with cross-linking version of mavacamten were entirely in the IHM state (16). Given this evidence, we predicted that a saturating dose of mavacamten should increase the FRET efficiency from 43% to 74%, if the IHM is fully populated, as predicted by the EM studies.

Mavacamten was added to our FRET-labeled sample, and FRET did indeed increase, but only by about 3% (Fig. 3A), an order of magnitude less than predicted if the IHM were fully populated. Increasing mavacamten resulted in a hyperbolic response with a B_max_ of 4.3% and K_d_ of 27 μM. The 3% FRET change fits to a model with a 4% increase in the population participating in FRET. The labeled myosin gave the same robust response to mavacamten (33% increase in SRX) as unlabeled myosin in the fluorescent nucleotide turnover kinetics studies (Fig. 3B).

**Figure 3.**
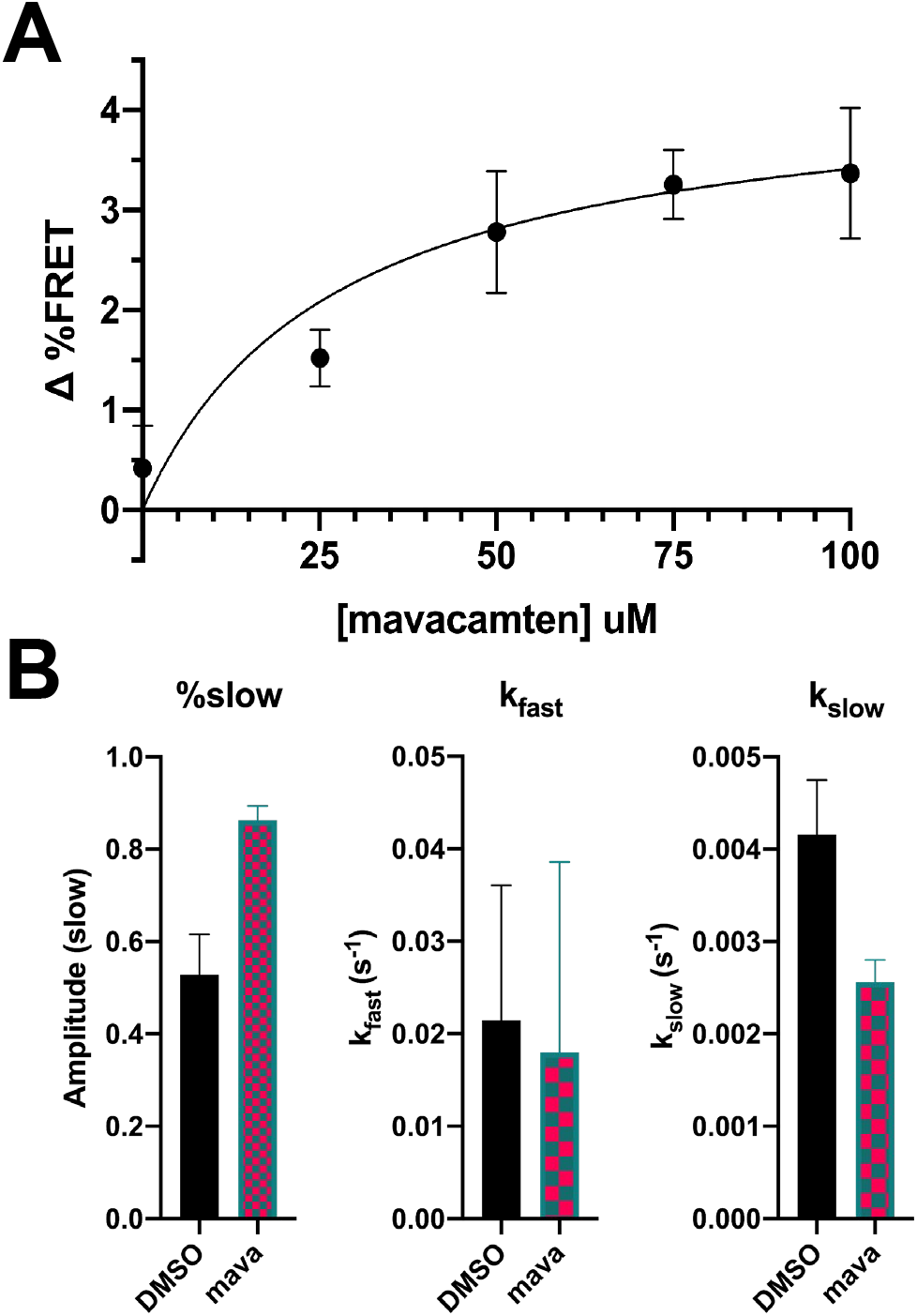
Mavacamten causes a small increase in FRET, but a large increase in SRX population in nucleotide turnover. (A) Mavacamten dosage-response curve of FRET of 5 μM labeled HMM (n=3 per condition) (B) Single nucleotide turnover of 0.2 μM labeled myosin with 10 μM mavacamten or 1% DMSO mixed with 4μM mant-ATP and chased with 2mM ATP (n=3 per condition)

## Discussion

Multiple forms of myosin, including mammalian cardiac myosin, appear to have a conserved mechanism of auto-inhibition (3). Here we have observed the IHM of beta-cardiac myosin, using TR-FRET with randomly distributed donor-labeled and acceptor-labeled RLC. We find that about 40% of the myosin molecules are in the IHM. The SRX measured by fluorescent nucleotide turnover has been measured to vary from 40% to 70% in different beta-cardiac myosin of various species (15–17). The comparable IHM and SRX populations suggest that the structurally defined and chemically defined auto-inhibited states of myosin are related.

The predicted distance in most reported models of the interacting heads motif of β-cardiac myosin, which are based on cryo-EM images of tarantula thick filaments (4,13,18), is around 5.0 nm, which is longer than the center distance seen by TR-FRET. In those EM studies, the region of lowest resolution is that of the RLC, suggesting that this area contains more disorder than the rest of the structure, so this discrepancy is unsurprising (18).

We were unable to observe the open conformation of myosin with this FRET system, indicating that most of the heads on these myosin molecules spend most of their time at least 4 nm apart, as predicted by most models.

We tested whether the shorter distance corresponds to the IHM by treating myosin with glutaraldehyde, a cross linker used to trap the IHM structure in myosin dimers for electron microscopy (3). We found that glutaraldehyde greatly increased FRET between donor- and acceptor-labeled RLC, confirming the IHM assignment.

Given the effectiveness of mavacamten to inhibit beta-cardiac myosin contractility (19) and help patients with hypertrophic cardiomyopathy (20), we hypothesized that mavacamten protects the heart through sequestering of myosin heads in the inactive SRX state, which is outside of the mechano-chemical cycle. Our lab and others have found that mavacamten potently increases the mole fraction of the SRX population (measured by nucleotide exchange) of cardiac myosin (15,16). However, we observed that mavacamten did not greatly increase the mole ratio of IHM (measured structurally by FRET). Since our study detects the structural state of IHM under conditions (no freezing, no fixing) similar to those used previously to detect SRX, these results call into question the previously untested hypothesis that there is a one-to-one correspondence between the IHM structural state and the SRX biochemical state.

Based on our results, we hypothesize that mavacamten does not fully trap cardiac myosin in the IHM. This may explain why it successfully treats hypertrophic cardiomyopathy without completely stopping heart contractions. Studies in cardiac S1, a cleaved single-headed version of myosin, showed that mavacamten primarily inhibits myosin by slowing the phosphate release step and ADP release in the mechano-chemical cycle, without head-head interaction (19). The slowing of these rates could also slow the single-turnover kinetics, leading to SRX-like rates. Our results suggest that the IHM-like structure leads to SRX-like kinetics, but that the population of myosin in SRX can greatly exceed that in the IHM. These SRX myosin heads have slower kinetics, sufficiently to be cardioprotective, but does not require static sequestration via the IHM structure.

## Experimental procedures

### Protein preparations

β-cardiac myosin was purified from bovine hearts obtained on wet ice from *Pel-Freez*, as described in Rohde *et al* (21). This isolation takes advantage of the tendency of myosin thick filaments to associate and dissociate in buffers of various ionic strengths, and cycling through multiple times leaves a pure sample of myosin. The entire purification process was performed at 4 °C.

Unlabeled β-cardiac HMM was prepared as described in our previous work (21). Full-length bovine β-cardiac myosin was digested to HMM in 10 mM tris(hydroxymethyl)aminomethane (Tris), 600 mM KCl, 2 mM MgCl_2_, and 1 mM DTT (pH 7.5) with α-chymotrypsin (Sigma-Aldrich, 0.025 mg/ml final concentration) for 10 minutes at 25 °C, followed by addition of pefabloc (Roche, 5 mM final concentration), then dialyzed into 10 mM tris(hydroxymethyl)aminomethane (Tris) (pH 7.5) with 2 mM MgCl_2_, followed by Q-sephadex ion-exchange chromatography purification. Intact HMM without contaminants as evaluated by SDS-Page were pooled for experiments.

Bovine cardiac RLC with a single reactive cysteine at position 105 was expressed in *Escherichia coli*, and purified by inclusion body isolation followed by ion exchange chromatography as described in our previous work (21). We labeled the purified RLC with 10 molar excess of 5-((((2-Iodoacetyl)amino)ethyl)amino)naphthalene-1-sulfonic acid (IAEDANS, Invitrogen) or dabcyl C2 maleimide (AnaSpec) via a maleimide-sulfhydryl reaction overnight at 4 °C and then removed free dye by gel filtration chromatography. The labeled RLC was snap frozen in liquid nitrogen and stored at −80 °C until use.

Labeling efficiency was determined by dye absorbance and Bradford protein concentration assay, as well as liquid chromatography mass spectroscopy. Labeling was confirmed using ultra-performance liquid chromatography-electrospray ionization mass spectrometry (UPLC-ESI MS). Unlabeled and fluorescently labeled RLC samples (≈50 mM) were buffer-exchanged into 10 mM ammonium bicarbonate using Zeba desalting columns according to manufacturer’s protocol (ThermoFisher Scientific). The sample was then run through a Waters C4 reversed phase chromatography column with a gradient of water (0.1% formic acid) and acetonitrile (0.1% formic acid) (low organic to high organic over 26 min linear gradient), 0.4 mL/min into the Synapt G2 QTOF MS which is scanned continuously from m/z 400 to m/z 2500.

Labeled β-cardiac HMM was prepared by first exchanging labeled RLC to full-length myosin and then enzymatically digesting to HMM. Full-length myosin was mixed with 4 times excess labeled RLC in 1:3 EDANS-dabcyl (or unlabeled for the donor-only sample) ratio and 12mM ethylenediaminetetraacetic acid (EDTA) and incubated for 10 minutes at 30°C, causing the endogenous RLC to dissociate from myosin. The addition of 14mM MgCl_2_ induces association of labeled RLC back on myosin. After the RLC exchange, the labeled myosin was brought to 2 mM MgCl_2_ in its freezing buffer before digestion to HMM with α-chymotrypsin (Sigma-Aldrich, 0.025 mg/ml final concentration) for 10 minutes at 25 °C, followed by addition of pefabloc (Roche, 5 mM final concentration) and then dialyzed into 10 mM Tris pH 7.5 with 2 mM MgCl2. Digested HMM was purified by ion-exchange chromatography (GE HiPrep) and confirmed by SDS-PAGE prior to experiments.

### Time-resolved FRET measurements

TR-FRET waveforms were measured by time-correlated single-photon counting after excitation with a 381 nm subnanosecond pulsed diode laser (LDH-P-C-375B, PicoQuant – Berlin, Germany). Emitted light was selected using a 440 ± 20 nm filter (Semrock, NY) and detected with a PMH-100 photomultiplier (Becker-Hickl). 10 cycles of photon counting were measured and averaged per sample, and at least 3 biochemically distinct samples were tested. Mavacamten was added at the indicated concentrations, along with 2mM magnesium ATP, and incubated for 10 minutes. Glutaraldehyde-treated samples (5 μM) in 10mM MOPS, 30mM KCl, 2mM MgATP, and 1mM DTT, pH 7.5 were treated with a 0.1% glutaraldehyde solution for 1 minute, then quenched with 100mM Tris (pH 7.5). The instrument response function (IRF) was recorded from water.

TR-FRET data was then analyzed by fitting fluorescence decay waveforms by non-linear regression analysis to a multi-exponential decay, as described in our previous work (21). The fluorescence decay can generally be described by a sum of exponentials,

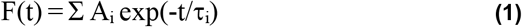

where τ_i_ is the fluorescence lifetime of the i-th component. Glutaraldehyde treatment produced fluorescent signals with low intensity and short lifetimes, in both labeled and unlabeled samples, so the glutaraldehyde-treated waveforms were analyzed with the donor-only sample treated with glutaraldehyde. The number of donor-only lifetimes τ_i_ was determined first by fitting F_D_(t) to Eq. 1, varying n from 1 to 4, choosing n as the smallest value needed to minimize χ2. The sample containing both donor and acceptor was fit to different structural models with the free parameters of mole ratio, and center distance and FWHM of one (or more) Gaussian-distributed population, and then the χ2 of the different models were compared, and the model that sufficiently minimized χ2 was chosen. The donor-acceptor waveform was also fit to the following model where mole fraction of the population participating in FRET is allowed to vary:

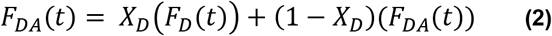

Where the mole fraction of donors participating in FRET is

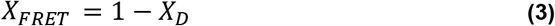

### Steady-state ATPase assay

The actin-activated MgATPase activity of purified cardiac myosin HMM or S1 was measured using an NADH-coupled assay performed at 25 °C in 10 mM Tris and 2 mM MgCl2 with 1 mM DTT (pH 7.5) in a 96-well clear plate. The reaction mix contained 0.2 μM HMM, varied actin concentrations, 0.2 mM NADH, 0.5 mM phosphoenol pyruvate, 2.1 mM ATP, 10 U/mL lactic acid dehydrogenase, and 40 U/mL pyruvate kinase. The conversion of NADH to NAD+ was measured on a SpectraMax 384 by monitoring absorbance at 340nm wavelength for 30 minutes. The activity was calculated from the slope obtained by linear regression, and divided by the concentration of catalytic sites in sample.

### Basal ATPase assay

Single-ATP turnover experiments with 2’- (or-3’)- O- (N-methylanthraniloyl) adenosine 5’-triphosphate (mant-ATP) were performed at 25 °C in 10 mM Tris, 25 mM potassium acetate, 4 mM MgCl2, 1 mM EDTA, and 1 mM DTT (pH 7.5). Steady-state fluorescence (total fluorescence intensity) was detected on an Applied Photophysics stopped-flow spectrophotometer capable of single-mix and sequential-mix experiments with water-bath temperature control. Samples were excited at 280 nm with a xenon lamp and monochromator and were detected through a 400-nm long-pass filter. The single-mix dead time for this instrument is 1.3 ms. All buffers were filtered and then degassed for 30 min under high vacuum before use.

### Actin-myosin binding assay

Rabbit skeletal actin labeled with pyrene (0.5μM) was mixed with HMM (labeled or unlabeled each at 1, 0.67, 0.5, 0.375, 0.28, 0.21, and 0.16μM) in F-buffer (10 mM Tris pH 7.5, 3 mM MgCl_2_). Pyrene fluorescence (Ex= 365nm, Em= 420nm) was measured in a 96-well clear plate by a Molecular Devices Gemini EM microplate spectrofluorometer. 2 mM ATP was added for actin dissociation, and pyrene fluorescence was measured again.

## Data availability

Raw data shared upon request to corresponding author, DDT.

## Acknowledgements

Fluorescence experiments were performed at the Biophysical Technology Center, University of Minnesota. We thank the Mass Spectroscopy Laboratory, University of Minnesota, for equipment and technical assistance. We thank Megan McCarthy, Lien Phung, and John Rohde for helpful discussions. We thank Destiny Ziebol for administrative assistance.

## Funding and additional information

This work was supported by NIH grants to DDT (R01AR032961, R37AG26160). SC was supported by NIH training grant T32AR007612.

## References

1. Sweeney, H. L., and Houdusse, A. (2010) Structural and functional insights into the Myosin motor mechanism. Annu Rev Biophys 39, 539–557

2. Woodhead, J. L., Zhao, F. Q., Craig, R., Egelman, E. H., Alamo, L., and Padron, R. (2005) Atomic model of a myosin filament in the relaxed state. Nature 436, 1195–1199

3. Lee, K. H., Sulbaran, G., Yang, S., Mun, J. Y., Alamo, L., Pinto, A., Sato, O., Ikebe, M., Liu, X., Korn, E. D., Sarsoza, F., Bernstein, S. I., Padron, R., and Craig, R. (2018) Interacting-heads motif has been conserved as a mechanism of myosin II inhibition since before the origin of animals. Proc Natl Acad Sci U S A 115, E1991–E2000

4. Alamo, L., Pinto, A., Sulbaran, G., Mavarez, J., and Padron, R. (2018) Lessons from a tarantula: new insights into myosin interacting-heads motif evolution and its implications on disease. Biophys Rev 10, 1465–1477

5. Jung, H. S., Komatsu, S., Ikebe, M., and Craig, R. (2008) Head-head and head-tail interaction: a general mechanism for switching off myosin II activity in cells. Mol Biol Cell 19, 3234–3242

6. Zoghbi, M. E., Woodhead, J. L., Moss, R. L., and Craig, R. (2008) Three-dimensional structure of vertebrate cardiac muscle myosin filaments. Proc Natl Acad Sci U S A 105, 2386–2390

7. Xu, J. Q., Harder, B. A., Uman, P., and Craig, R. (1996) Myosin filament structure in vertebrate smooth muscle. J Cell Biol 134, 53–66

8. Linari, M., Brunello, E., Reconditi, M., Fusi, L., Caremani, M., Narayanan, T., Piazzesi, G., Lombardi, V., and Irving, M. (2015) Force generation by skeletal muscle is controlled by mechanosensing in myosin filaments. Nature 528, 276–279

9. Caremani, M., Pinzauti, F., Powers, J. D., Governali, S., Narayanan, T., Stienen, G. J. M., Reconditi, M., Linari, M., Lombardi, V., and Piazzesi, G. (2019) Inotropic interventions do not change the resting state of myosin motors during cardiac diastole. J Gen Physiol 151, 53–65

10. Fusi, L., Brunello, E., Yan, Z., and Irving, M. (2016) Thick filament mechano-sensing is a calcium-independent regulatory mechanism in skeletal muscle. Nat Commun 7, 13281

11. Brunello, E., Fusi, L., Ghisleni, A., Park-Holohan, S. J., Ovejero, J. G., Narayanan, T., and Irving, M. (2020) Myosin filament-based regulation of the dynamics of contraction in heart muscle. Proc Natl Acad Sci U S A 117, 8177–8186

12. Woodhead, J. L., and Craig, R. (2020) The mesa trail and the interacting heads motif of myosin II. Arch Biochem Biophys 680, 108228

13. Nag, S., Trivedi, D. V., Sarkar, S. S., Adhikari, A. S., Sunitha, M. S., Sutton, S., Ruppel, K. M., and Spudich, J. A. (2017) The myosin mesa and the basis of hypercontractility caused by hypertrophic cardiomyopathy mutations. Nat Struct Mol Biol 24, 525–533

14. Hooijman, P., Stewart, M. A., and Cooke, R. (2011) A new state of cardiac myosin with very slow ATP turnover: a potential cardioprotective mechanism in the heart. Biophys J 100, 1969–1976

15. Rohde, J. A., Roopnarine, O., Thomas, D. D., and Muretta, J. M. (2018) Mavacamten stabilizes an autoinhibited state of two-headed cardiac myosin. Proc Natl Acad Sci U S A 115, E7486–E7494

16. Anderson, R. L., Trivedi, D. V., Sarkar, S. S., Henze, M., Ma, W., Gong, H., Rogers, C. S., Gorham, J. M., Wong, F. L., Morck, M. M., Seidman, J. G., Ruppel, K. M., Irving, T. C., Cooke, R., Green, E. M., and Spudich, J. A. (2018) Deciphering the super relaxed state of human beta-cardiac myosin and the mode of action of mavacamten from myosin molecules to muscle fibers. Proc Natl Acad Sci U S A 115, E8143–E8152

17. Toepfer, C. N., Garfinkel, A. C., Venturini, G., Wakimoto, H., Repetti, G., Alamo, L., Sharma, A., Agarwal, R., Ewoldt, J. F., Cloonan, P., Letendre, J., Lun, M., Olivotto, I., Colan, S., Ashley, E., Jacoby, D., Michels, M., Redwood, C. S., Watkins, H. C., Day, S. M., Staples, J. F., Padron, R., Chopra, A., Ho, C. Y., Chen, C. S., Pereira, A. C., Seidman, J. G., and Seidman, C. E. (2020) Myosin Sequestration Regulates Sarcomere Function, Cardiomyocyte Energetics, and Metabolism, Informing the Pathogenesis of Hypertrophic Cardiomyopathy. Circulation 141, 828–842

18. Robert-Paganin, J., Auguin, D., and Houdusse, A. (2018) Hypertrophic cardiomyopathy disease results from disparate impairments of cardiac myosin function and auto-inhibition. Nat Commun 9, 4019

19. Kawas, R. F., Anderson, R. L., Ingle, S. R. B., Song, Y., Sran, A. S., and Rodriguez, H. M. (2017) A small-molecule modulator of cardiac myosin acts on multiple stages of the myosin chemomechanical cycle. J Biol Chem 292, 16571–16577

20. Olivotto, I., Oreziak, A., Barriales-Villa, R., Abraham, T. P., Masri, A., Garcia-Pavia, P., Saberi, S., Lakdawala, N. K., Wheeler, M. T., Owens, A., Kubanek, M., Wojakowski, W., Jensen, M. K., Gimeno-Blanes, J., Afshar, K., Myers, J., Hegde, S. M., Solomon, S. D., Sehnert, A. J., Zhang, D., Li, W., Bhattacharya, M., Edelberg, J. M., Waldman, C. B., Lester, S. J., Wang, A., Ho, C. Y., Jacoby, D., and investigators, E.-H. s. (2020) Mavacamten for treatment of symptomatic obstructive hypertrophic cardiomyopathy (EXPLORER-HCM): a randomised, double-blind, placebo-controlled, phase 3 trial. Lancet 396, 759–769

21. Rohde, J. A., Thomas, D. D., and Muretta, J. M. (2017) Heart failure drug changes the mechanoenzymology of the cardiac myosin powerstroke. Proc Natl Acad Sci U S A 114, E1796–E1804

